# The Energy Barrier Model in Membrane Biophysics: Ion Flow, Current-Voltage Relations, and Donnan Osmosis

**DOI:** 10.1101/2025.08.20.671285

**Authors:** Gerald S. Manning

## Abstract

An energy barrier model for a membrane yields new biophysical results. Here we explore both the equilibrium and time-dependent Nernst potential, capacitance and resistance, the resting potential, and osmosis.

## 1 Introduction

Biological membranes are heterogeneous assemblies, and their penetrability by water and ions reflects the complexity of their structure [1–4]. Water molecules are sparingly soluble in the lipid bilayer and can cross it directly or through hydrophilic protein channels. Small ions like sodium, potassium, and chloride are insoluble in the lipid but can be ferried across by soluble carriers or flow through ion selective channels. There are specialized protein channels for each of these ubiquitous ions. Notwithstanding the inevitably complicated nature of any biological system on a molecular level, simple models that capture some essential features may provide insights and suggest areas for measurements or simulations [5–7]. To enter and traverse a membrane a water molecule or an ion must acquire sufficient energy, so an exploration of an energy barrier model seems worth-while.

A calculation of the osmotic pressure of a simple binary electrolyte will serve to illustrate the central narrative of this essay. Suppose the (*y, z*) plane at *x* = 0 is considered as an ideal semipermeable membrane (see Figure 1–all figures are located at the end). The space *x* < 0 is occupied by pure water, and is designated as the “outer” compartment. The space *x* > 0 is occupied by an aqueous solution of a fully dissociated uni:univalent salt with a “+” cation and a “−” anion. This space is the “inner” compartment. The membrane is invisible to water molecules, which pass freely between outer and inner compartments, that is, across *x* = 0. The membrane is also invisible to the + cations, which are therefore said to be freely permeant to the membrane. However, it presents an infinitely high energy barrier to the “−” anions, which are then called impermeant.

**Figure 1.**
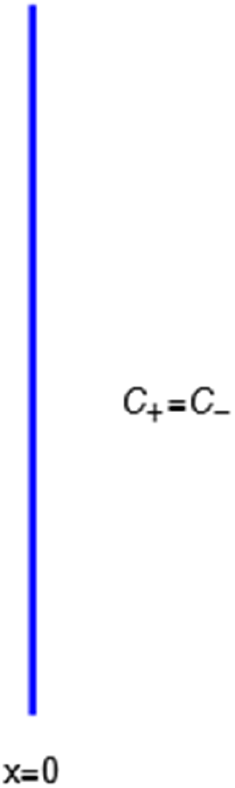
The energy barrier is the (y,z) plane seen end-on at x=0. The x coordinate increases from left to right. An electroneutral aqueous binary salt solution is in the half-volume x>0. The half-volume x<0 contains pure water. The barrier is invisible to the water but impermeable to the anion. The cation is freely permeant.

Consider the ionic fluxes *j*_+_ and *j*_−_ across the membrane at *x* = 0, that is, the number of + or − ions crossing the (*y, z*) plane at *x* = 0 per unit area per second. For the anion,

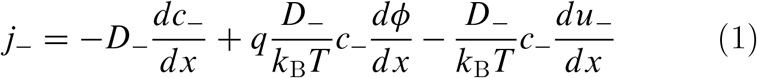

In this equation *c*_−_ is the concentration of − ions (number of ions per unit volume), *D*_−_ is a diffusion coefficient, *k*_B_*T* is the product of the Boltzmann constant and Kelvin temperature, *q* is the unit positive charge (the charge on the univalent cation is +*q*, and on the univalent anion it is −*q*), *ϕ* is the electrostatic potential, and *u*_−_ is the nonelectrical energy height that the anion must surmount to cross the membrane. The equation uses Einstein’s recognition that *D*_−_*/k*_B_*T* is the mobility of the ion. The anion is impermeant, which means first of all that *j*_−_ = 0. After canceling *D*_−_, the equation takes the form,

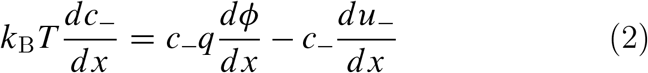

On the right side of this equation stands the total force per unit volume on the − ions, the sum of the electrical force (first term) and the mechanical force (second term) originating from the membrane material that prevents the impermeant anions from crossing the membrane.

Next we look at the flux of + ions, each bearing charge *q*,

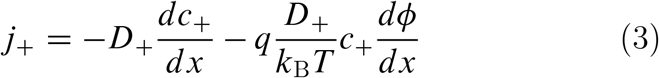

There is no term in this equation corresponding to the mechanical membrane-ion force, because the + ion is freely permeant, that is, *u*_+_ = 0. Nonetheless, the cation flux *j*_+_ must vanish along with the flux of impermeant anion because electroneutrality requires *c*_+_ = *c*_−_ everywhere. The equation for the cation corresponding to eq ?? for the anion is therefore,

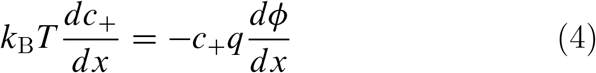

The force per unit volume on the + ions stems only from an electric field which prevents these ions from leaving the impermeant anions behind.

To complete the derivation of osmotic pressure, set *c*_+_ and *c*_−_ both equal to the salt concentration *c*_*s*_, and add the equations for cation and anion, eqs ?? and ??,

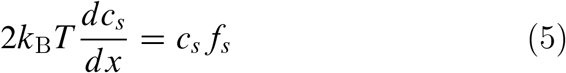

where *c*_*s*_*f*_*s*_ = −*c*_−_*du*_−_/*dx*, the electrical forces having canceled. The right-hand side is just the force per unit volume on a volume element at *x* (the direct forces repelling the impermeant anions from the membrane are transmitted to all the water molecules and cations in the volume element by collisional kinetic energy, i.e., by temperature). At equilibrium this volume force is balanced by a pressure gradient, *c*_*s*_*f*_*s*_ = *dP*/*dx*, and equating the two expressions for *c*_*s*_*f*_*s*_ shows that *dP*/*dx* = *2k*_B_*Tdc*_*s*_/*dx*. Since the salt concentration in the outer compartment is zero, integration across the membrane gives the van’t Hoff equation for the equilibrium pressure difference (osmotic pressure).

The factor 2 is frequently explained qualitatively by noting that the solute concentration in van’t Hoff’s equation is the total concentration of osmotically active particles, or “osmolarity”, in this case the total concentration of ions, cation plus anion. It also emerges automatically from the rigorous thermodynamic derivation involving the chemical potential of the solvent water. The insight gained here is that *2k*_B_*Tc*_*s*_ is actually the sum of two terms which have distinct physical origins. If the anion is impermeant, there is a term *k*_B_*Tc*_−_ stemming from a mechanical repulsion from the membrane, while even if the cation is mechanically freely permeant, it contributes a term *k*_B_*Tc*_+_ from the repulsive electric field arising from electroneutrality.

The electric field that arises from electroneutrality is present in eq ??. This equation should be familiar to neurophysiologists and biophysicists, for it is the differential form of the Nernst potential that plays such a central role in those fields. Division of this equation by *c*_+_ and integration across *x* = 0 gives the familiar expression for the Nernst potential of the + ion,

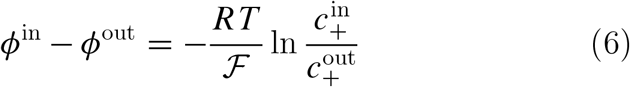

where Avogadro’s number has been used to write *RT*/ ℱ where *R* is the gas constant and ℱ = *N*_Av_*q* is the Faraday. If one wishes, *ϕ*^out^ may be set equal to zero. For the case under discussion 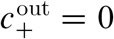, so the Nernst potential is −∞, signifying that the + ion cannot leave the salt solution in the inner compartment. Here and subsequently, mathematical liberties for the sake of physical transparency are deliberate.

The interconnectedness of electrical and mechanical forces when charged ions are involved begins to become apparent from these considerations. In many of the subsequent calculations, a quantity *a*_*i*_ = exp(−*u*_*i*_*/k*_B_*T*) is involved, where *u*_*i*_ is the mechanical energy impeding ions of species *i* from entering the membrane. Although this quantity has the form of a partition coefficient, it should not be so mistaken. In the above example, *u*_+_ is zero, indicating that were it not for its electrical charge, a + ion would be freely permeant and be present in the membrane at the same concentration as in free solution. Yet it has been rendered effectively impermeant because electroneutrality requires its presence to compensate the negative charge of the mechanically impermeant anions. Eq ?? shows that the force preventing the + ion from passing across the membrane is purely electrical.

In fact it arises entirely from the Nernst potential.

The use of a mechanical energy *u*_*i*_ to indicate the work that would have to be done on an ion *i*, if it lacked its electrical charge, to place it from free solution into the interior of a membrane allows a sharp definition of the meaning of “impermeant” and “permeant”. An ion *i* is impermeant if *u*_*i*_ = +∞. It is “freely permeant” if *u*_*i*_ = 0. It is permeant, but not freely so, if *u*_*i*_ is greater than zero but not infinite. (Negative values would indicate absorption by the membrane but this situation is not treated here.)

In the previous example the energy barrier is localized to the zero-width (*y, z*) plane. A minimal model for all subsequent purposes represents a membrane of finite width *h* by the mechanical energy barrier shown in Figure 2. The mechanical energy *u* of a molecule or ion equals zero in the outer compartment *x* < 0 and also equals zero in the inner compartment *x* > 0. It equals the constant value *u* in the space 0 < *x* < *h* occupied by the membrane. The mechanical force −*du/dx* exerted by the membrane material on an ion in the interior of the membrane 0 < *x* < h is zero, since *u* is constant there. At the surfaces *x* = 0 and *x* = *h, du/dx* is infinite but has clear physical meaning. If at *x* = 0, for example, *du/dx* is integrated from just outside the membrane where *u* = 0 to just inside where *u* = *ū*, the value of the integral is *ū*. On the other side of the membrane at *x* = *h*, if *du/dx* is integrated from just inside the membrane to just outside (left to right in Figure 2, in the direction of increasing *x*), the value of the integral is −*ū*. The notation can be simplified by omitting the bar on *u* when the meaning is clear.

**Figure 2.**
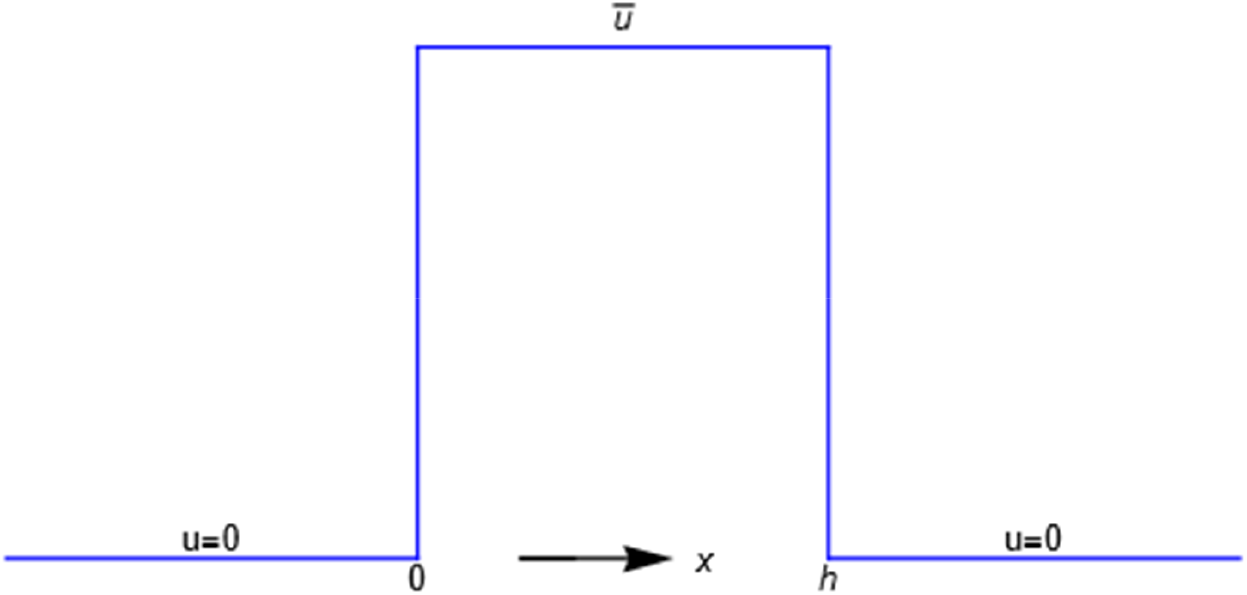
An energy barrier of width h. A particle has zero mechanical energy u in compartments x<0 and x>h but work u>0 must be done on it for entrance into the membrane 0<x<h. The energy acquired is shed as the particle exits the membrane.

To the author’s awareness, the first indication that an energy barrier can insightfully model interesting aspects of membrane physics was the application by Peter Debye to osmotic pressure [8, 9]. An extension of the Debye model to osmosis [10, 11] and to a statistical mechanical interpretation of the Kedem-Katchalsky thermodynamic flow equations followed [10]. Davson and Danielli [1] attempted to design details of energy barrier profiles that would describe what they knew or suspected of the molecular-level interaction of biological membranes with water and ions, but their thinking was in the pre-computational era. Eyring and others applied rate theory to an energy barrier model for membranes [2]. The minimal energy barrier model used here is intended to provide information on a coarse-grained level that might elude easy interpretation of computational results at molecular- and atomic-level resolution.

Since in the interior of this minimal energy barrier model, ions and water molecules move as they would in a macroscopic fluid phase, the first topic to be considered is free diffusion of the ions of a binary electrolyte like KCl or NaCl [12, 13]. Emphasis is placed on the electrostatic diffusion potential [14] and its physical origins [15]. The concept of polarization charge density from the classical theory of electricity in polarizable media is introduced [15–17]. The next subject is the Nernst potential [3,4] and its structure across an energy barrier. A current-voltage analysis for the energy barrier follows. It recaptures the differential equation for the equivalent ℛ 𝒞 electrical circuit of neurophysiology [3] but is based on a molecular-level interpretation of both the resistance ℛ and capacitance 𝒞.

Since an ionic system of great interest in biology involves the simultaneous presence of sodium, potassium, and chloride ions, free diffusion in this example of a solution of mixed salts becomes an objective of theoretical study. A straightforward generalization of the binary salt case is not possible [12, 13, 18], and a careful inspection of the linear and nonlinear properties of the Nernst-Plank equations is involved. The result is required for the calculation of the resting potential [3] in the energy barrier model, where, however, the treatment of the discontinuities at the membrane-solution surfaces turns out to be essential. Attention is then directed to Donnan osmosis, that is, the flow of water through the membrane in the presence of both impermeant and permeant ions. Although all of these questions have been much discussed, the present analysis proceeds differently, recapturing some familiar results hopefully in an interesting way but also arriving at some very different ones.

## II. FREE DIFFUSION OF A BINARY ELECTROLYTE

A simple binary electrolyte (or salt) like NaCl or KCl diffuses in free solution as an electrically neutral component even though its constituent ions are completely dissociated and move as separate entities. The well-known Nernst-Hartley equation

[13] expresses the diffusion constant of the neutral salt in terms of the diffusion constants, or mobilities, of the individual ions. Both the diffusion constants of a large variety of salts and the mobilities of their ions have been accurately measured [13] and found to be in close agreement with the Nernst-Hartley theory, even when the salt contains divalent or trivalent ions [15].

At the core of the Nernst-Hartley theory is the assumption that electroneutrality holds in volume elements within the diffusing concentration gradient. At the same time, however, the assumption that a self-generated electrical field exists within the concentration gradient is also central. An equation for this diffusion potential is also provided by the theory which shows that the field arises from the differing mobilities of the cations and anions. Unlike the diffusion constant of the salt and the mobilities of the ions, the measurement of the diffusion potential is in principle difficult and has been regarded with unease. Can an electric field, not applied externally, be present in an electroneutral volume element? In this section we review the Nernst-Hartley theory and show how this question has been resolved [15] with no recourse to a fictitious space charge, which has been rejected in the physical chemistry of electrolyte solutions [13].

With the restriction for simplicity to 1:1 electrolytes, the diffusion fluxes of cation and anion are,

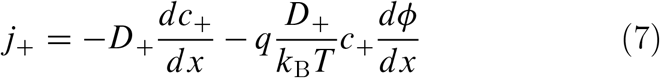

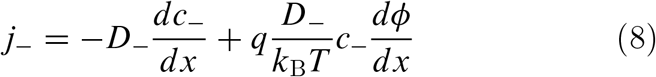

Electroneutrality dictates that *c*_+_ = *c*_−_, and *c*_*s*_ then stands for both concentrations. Electroneutrality and the consequent absence of space charge during diffusion also requires that *j*_+_ = *j*_−_, and *j*_*s*_ stands for both fluxes. Divide each of these equations by their corresponding diffusion constants, and then add the two equations. The gradient of electrostatic potential *ϕ* cancels, and the resulting equation can be solved for the salt flux *j*_*s*_, yielding the Nernst-Hartley relation,

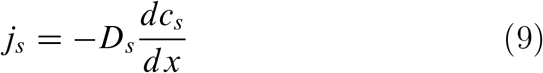

where,

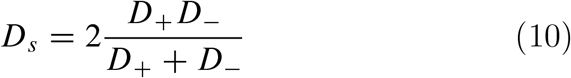

As seen, the salt flux is just Fick’s law for diffusion of the salt with the diffusion constant composed of the diffusion constants of the individual ions.

To get the diffusion potential, subtract the two ion flux equations, remembering that *j*_+_ and *j*_−_ are equal and therefore cancel. Solve for *dϕ/dx*,

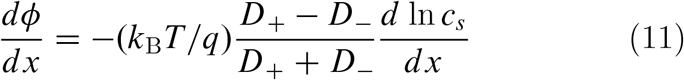

The electric field is the negative of *dϕ/dx*. The equation shows that it arises from the difference in diffusion constants (mobilities through the Einstein relation) of the cations and anions. The diffusion potential has the same theoretical basis as the Nernst-Hartley diffusion flux, which has been thoroughly verified by direct measurements [15].

The next step takes the Nernst-Hartley theory a bit further [15]. Differentiate *dϕ/dx* once again with respect to *x*,

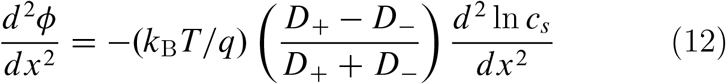

But the fundamental form of the Poisson equation from the theory of electric fields is [16,17],

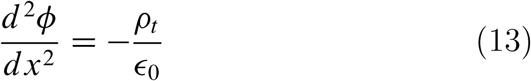

where *ρ*_*t*_ is the total charge density, and *ϵ* _0_ is the permittivity of vacuum. In a polarizable medium the total charge density consists of two parts, *ρ*_*t*_ = *ρ* + *ρ*_*P*_. The contribution *ρ* is from the space charge density. Given electroneutrality, it equals zero in salt diffusion. The contribution *ρ* _*P*_ is called the polarization charge density. Its origin is from the dipoles, permanent or induced, as aligned by the field −*dϕ/dx*. In the diffusion of a binary salt, the cations and anions have different mobilities, and the net charge density in a volume element, although zero by electroneutrality, is polarized. If the cations move more quickly than the anions, the positive ends of the charge distribution dipoles will be biased in the direction of lower salt concentration (that is, in the direction of diffusion velocity). An equation for the polarization charge density of a diffusing binary salt is obtained by equating the right-hand sides of the previous two equations with *ρ* _*t*_ = *ρ* _*P*_,

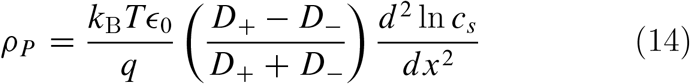

This equation has the same theoretical basis as the Nernst-Hartley diffusion equation.

The polarization charge density is not a familiar concept in biophysics, but it is an integral part of the theory of electrostatics of polarizable media and appears in all of the standard textbooks on the classical theory of electricity and magnetism. Cartoons exist as a visual aid [17], and Panofsky and Phillips [16] offer a verbal description along with their rigorous mathematical analysis, “If we have an inhomogeneous dipole moment per unit volume, *ρ*_*P*_ will represent the charge density that accumulates from incomplete cancellation of the ends of the individual dipoles distributed in the volume… [It] will vanish in a homogeneous medium.” In diffusion the medium is not homogeneous, since there is a nonvanishing concentration gradient.

## III. THE NERNST POTENTIAL AND THE POLARIZATION CHARGE DENSITY

The Nernst electrostatic potential has been mentioned in previous sections in various contexts. What is the Nernst potential? Suppose that in two distinct volume elements at *x*_1_ and *x*_2_ in an ordinary electrolyte solution like aqueous NaCl there is momentarily a difference in electrostatic potential *ϕ*(*x*_2_) − *ϕ*(*x*_1_), and that in these two volume elements the concentrations of one species of the ions comprising the electrolyte, say a “+” cation with unit positive charge *q*, happens momentarily to be distributed according to a Boltzmann equilibrium probability,

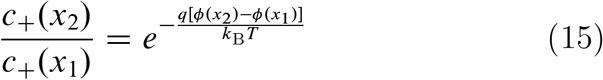

Taking logarithms and rearranging gives,

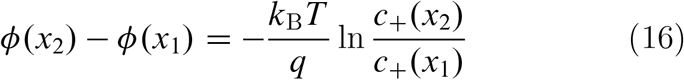

This potential difference, or voltage, is called the Nernst potential for a univalent cation. If the ion is a univalent anion with charge −*q*, the sign on the right-hand side is changed to a plus.

The importance of the Nernst potential goes well beyond a highly improbable simultaneous voltage and ionic concentration fluctuation. Suppose we have the more realistic situation of a membrane separating two solutions of the same 1:1 electrolyte but at different concentrations. Suppose one of the ions, say the anion, is impermeant to the membrane, while the cation is permeant. The Nernst potential then gives the voltage across the membrane at equilibrium, which can be achieved since the cations can exchange between the two sides of the membrane. The standard thermodynamic proof of this statement proceeds by equating the cation electrochemical potential (chemical potential of the cation supplemented by its electrostatic energy) in each of the two solutions, whereupon the Nernst potential immediately follows. A crucial attribute of a thermodynamic analysis is that it returns the correct answer. Another characteristic is that it may not provide much physical insight. Attempts at mechanistic interpretation based on charge separation are essentially the same as the incorrect assertion of charge separation in free diffusion of ions of differing mobilities (see Chapter 2). An energy barrier model may be helpful.

Consider an energy barrier that separates two uni-univalent salt solutions of different concentrations (see Figure 3). The “−” anion is impermeant, a word with sharp meaning as presently used; the energy *u*_−_ of the impermeant anion inside the energy barrier (the membrane) is equal to +∞. The + cation is permeant, meaning that its energy *u*_+_ inside the barrier is greater than zero but has some definite value less than ∞ (for a freely permeant ion, the value *u*_+_ would equal zero). Outside the barrier, in both solutions, the barrier energies of the ions are zero. The equation for the flux of the cation is,

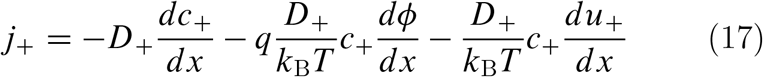

**Figure 3.**
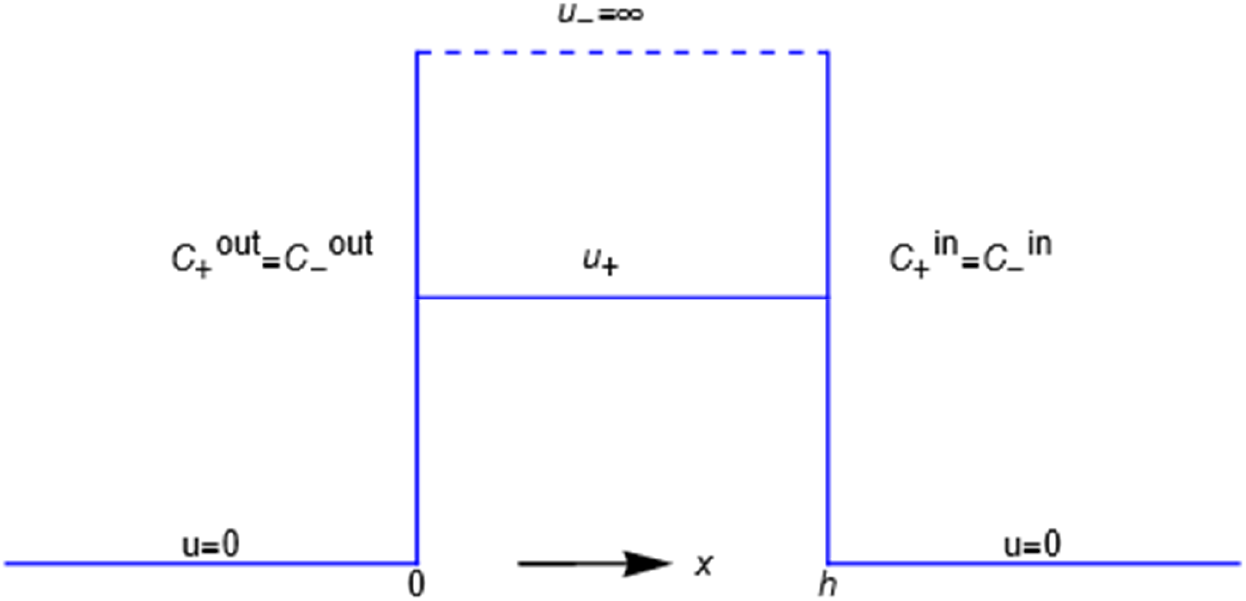
Profiles of the potential energies u_i_ representing the mechanical interactions of the membrane with the + and - ions of a binary salt of unequal but electroneutral concentrations in the outer and inner compartments.

Since the anion is impermeant, its flux *j*_−_=0. The electrical current is i= −*q(j*_+_ −*j*_−_) (the sign makes no difference for now, and is explained later). The Nernst potential pertains to a system with no net electrical current. Since i=0, and *j*_−_ = 0, it is also true that *j*_+_ = 0, and after canceling *D*_+_ and dividing by *c*_+_, this equation can be rearranged to,

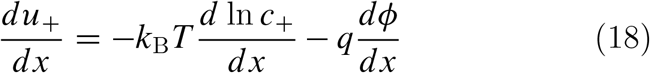

Now integrate the left side of the equation left to right from the “outer” solution to the “inner” one. Since *u*_+_ = 0 outside the membrane in both solutions, the integration results in zero. The corresponding integration of the right-hand side then gives the Nernst potential,

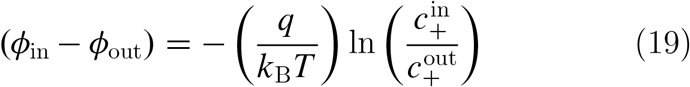

This derivation is almost the same as the one in the Introduction section, but it serves here to introduce a membrane of finite width and the necessary presence of an impermeant ion of opposite charge in addition to the ion associated with the Nernst potential.

The derivation also offers the possibility of additional information. Integrate eq. ?? across the left interface, in other words across the discontinuity at *x* = 0. To do so, notice that to the left of the discontinuity, there are the equalities 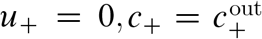, and *ϕ* = *ϕ*_out_. Just to the right of the discontinuity at *x* = 0, that is, just inside the membrane, *u*_+_ equals its constant value (which we also call *u*_+_) inside the energy barrier, *ϕ* is set equal to its value *ϕ*′ just inside the membrane at *x* = 0, and by electroneutrality, *c*′ = 0 inside the membrane, since the concentration of the impermeant anion is zero there. The result of the integration is,

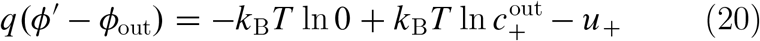

The last two terms on the right-hand side have finite values (in particular, the cation is permeant), but of course − ln 0 = +∞ and therefore dominates, so the potential jumps to infinity just inside the membrane. We find the same thing if eq. ?? is integrated across the discontinuity at *x* = *h*, namely, the value of the potential *ϕ* ^″^ just inside the membrane at *x* = *h* also jumps to +∞. The situation is drawn in Figure 4 for 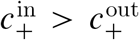, in other words, for a negative Nernst potential.

**Figure 4.**
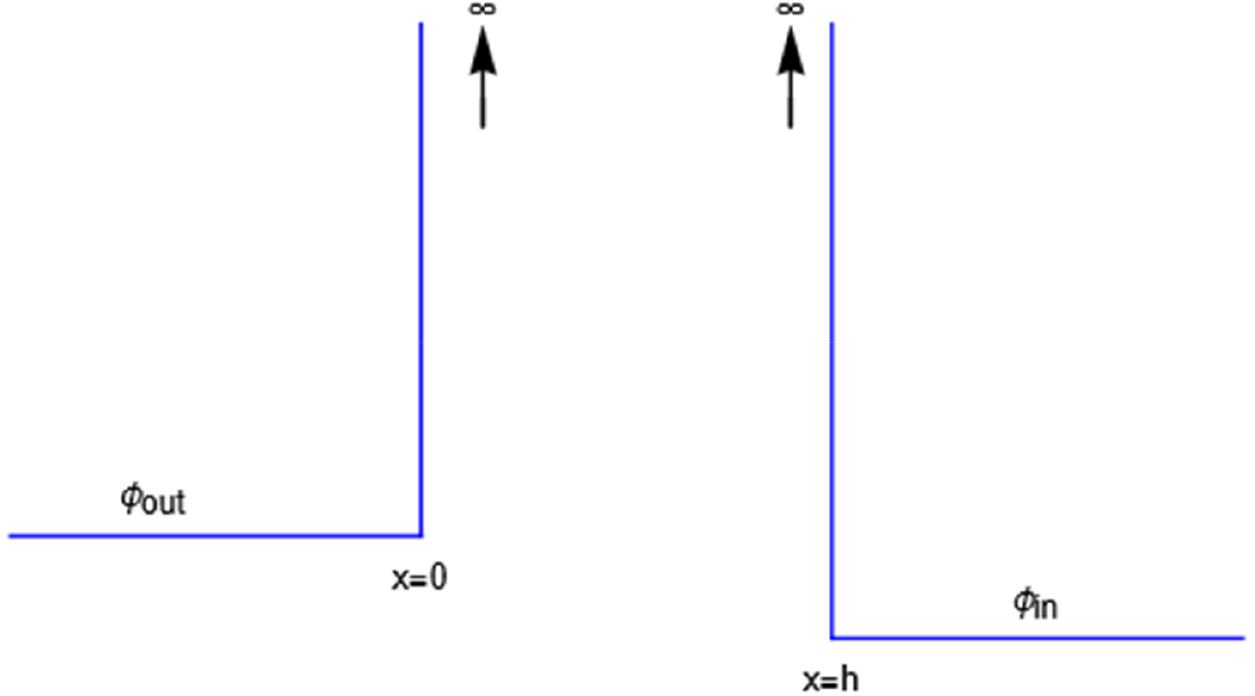
Illustration of the Nernst potential φ(x) for the + cation across an energy barrier of width h. As x approaches 0 from the left, φ(x) jumps to infinity from its uniform value φ_out_ in the outer compartment. As x exits the barrier at x=h, φ(x) drops from infinity to its value φ_in_ in the inner compartment. The Nernst potential is φ_in_-φ_out_, which is negative in this figure, where the concentration of the 1:1 binary salt, hence the cation, is greater in the inner compartment than in the outer. The anion is impermeant.

If the integration of eq. ?? is carried out from the left side of the energy barrier to the right side, remaining in the interior, the result is,

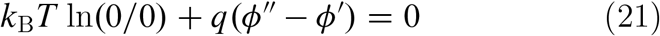

since *u*′ is constant throughout the interior, and *c*′ = 0 everywhere in the interior. A consistent physical interpretation of this otherwise indeterminant equation is that there is no electric field inside the energy barrier since there is zero net charge (electroneutrality) at either end. The potential is then constant throughout the interior, so *ϕ* ^″^ = *ϕ* ^′^. As for ln(0/0) = 0, it is just the statement that ln(*c*′*/c*′) = ln 1 = 0 for the constant vanishing value of *c*′ in the interior.

### The polarization charge density

The system under discussion in this chapter is similar to the case of free diffusion of a binary salt in the absence of a membrane, as discussed and documented in Chapter II. In free diffusion one of the ions, say the cation, has a mobility greater than the other. This ion then diffuses faster than the other, and a polarization of the overall ion distribution occurs with no actual separation of charges that would violate electroneutrality. The evidence for local electroneutrality inside the salt concentration gradient is strong. The Nernst-Hartley theory (not to be confused with the Nernst potential) gives a formula for the diffusion coefficient of the salt in terms of the diffusion coefficients (mobilities) of the individual ions. The theory is based on the assumption of local electroneutrality. The theory is in very close agreement with experimental measurements of both the salt diffusion coefficient and the individual mobilities.

When an impermeant anion encounters the surface of a membrane, it recoils from it, its mobility in the direction of the membrane is forced to reduce to zero, while the permeant cation can enter. The resulting polarization at the membrane surface does not require a violation of electroneutrality. There is no reason to believe that the cations separate from the anions and pass all the way across the membrane to the other side (of course there can be an equilibrating exchange of cations).

The calculation of the polarization charge density at the surfaces of the membrane is similar to that for free diffusion in Chapter II. It proceeds by differentiating eq ??,

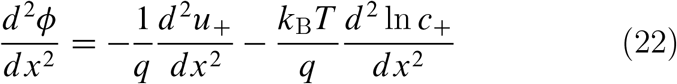

But from Poisson’s equation,

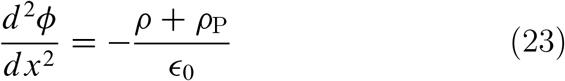

where the free charge density ρ = 0 by electroneutrality, leaving the polarization charge density *p*_P_. Therefore,

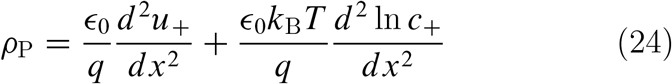

This equation is the same as for free diffusion but for the energy barrier *u*′ term. The second derivatives are zero everywhere except at the boundary discontinuities. They are mathematically not well-defined there, but the physical meaning, that the charge polarization takes place at the membrane surfaces, should be clear.

## IV. CAPACITANCE AND CURRENT-VOLTAGE RELATION

The system at hand is the same as in the previous chapter (see Figure 3), two 1:1 salt solutions of the same ions but different concentrations are separated by an energy barrier, impermeable to the anion but permeable to the cation. Suppose the system is initially at rest. We have seen in the previous chapter that the voltage across the energy barrier is then given by the Nernst potential. If a steady current is imposed at time *t* = 0, the voltage will change in time from the Nernst value to a value consistent with the applied current. In another situation the system initially may not be at rest but instead is in a steady state imposed by an applied current. If at *t* = 0 the current is then turned off, the voltage will relax in time to the Nernst value.

The time course of the voltage in these cases is determined by the capacitance of the energy barrier, and how to bring capacitance into the equations of electrodiffusion is the topic of this chapter.

The definition of capacitance 𝒞 is 𝒞 = *Q/V*, that is, 𝒞 is the charge *Q* per unit volt across the capacitor, where *Q* is the positive charge on one plate of the capacitor while −*Q* is the charge on the opposite parallel plate. But since *dQ/dt* is an electrical current *i, d V/dt* = *i/* 𝒞, and that is why in a simple electrical circuit, capacitance governs the time course of voltage. As for the origin of the charge *Q*, the charge flowing as current “piles up” [6] at one surface of the capacitor and is correspondingly depleted at the opposite surface.

The energy barrier model for a membrane provides a natural way to envision the piling up of charge in capacitor-like fashion. Recall the Nernst potential from the previous chapter in a current-free situation. There is a polarization charge density at each of the two surfaces of the energy barrier. When a current flows in response to an imposed electric field, the polarization charge density is itself further polarized. The field-induced perturbation of the equilibrium polarization charge density can be represented by charges *Q* and −*Q* at the surfaces of the energy barrier.

The flux of + cations is,

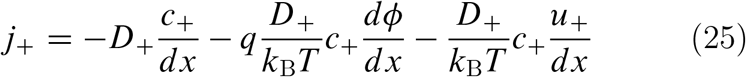

while *j*_−_ = 0 since the anion is impermeant. The electrical current *i* equals −*q*(*j*′ − *j*_−_) (we follow neurophysiological convention for direction of positive current opposite to direction of positive flux [3]). Since *j*_−_ = 0, the current is −*qj*′. The current is not zero in this chapter, but electroneutrality requires *c*′ = *c*_−_ in all volume elements.

From these equations and definitions, an expression for the current is,

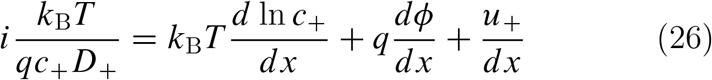

The goal here is to develop a molecular-level analog to a linear electric circuit, and to that end a linear mathematical environment is suitable. On the left-hand side of this equation, to retain the dependence of *c*′ on location *x* is to retain nonlinearity of the left-hand side, since the current *i* is represented as a linear flow (it appears to the first power). Therefore *c*′ on the lefthand side of the equation is taken as constant. The left side is then linear, along with the three linear force terms on the right side (including the concentration gradient as a virtual force).

The next step is to recognize that charge can “pile up” against the energy barrier (and be correspondingly depleted at the other side of the barrier) by representing the charge distribution *Q*(*x, t*) in space and time by the expression,

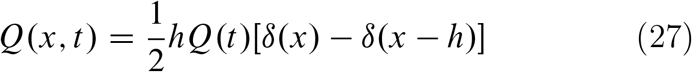

where the Dirac delta functions [17] on the right are sharply peaked at the two membrane surfaces *x* = 0 and *x* = *h* and equal zero elsewhere.

Integration of the current *i* across the left membrane surface at *x* = 0, left to right (see Figure 3), from outside the membrane to just inside, proceeds as follows:

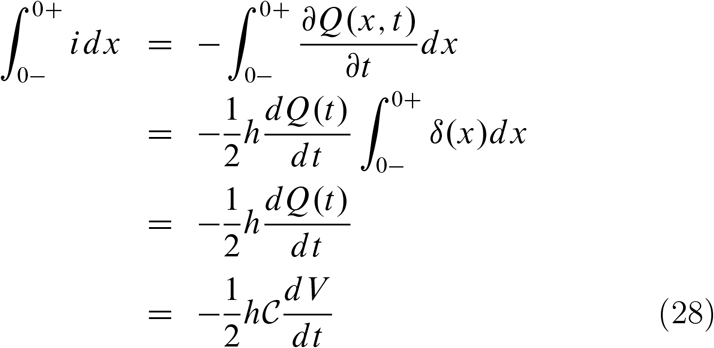

The minus sign in the first line conforms to the direction of current flow. Exactly the same result is obtained when the current is integrated across the membrane surface at *x* = *h*, since then the minus sign is evident from the charge distribution in eq ??,

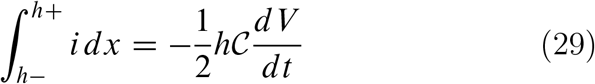

The next step is to use these results in integrating eq ??, first across the surface at *x* = 0, then across the surface at *x* = *h*. The two equations thus obtained are added (the addition cancels the *u*′ force terms) with the result,

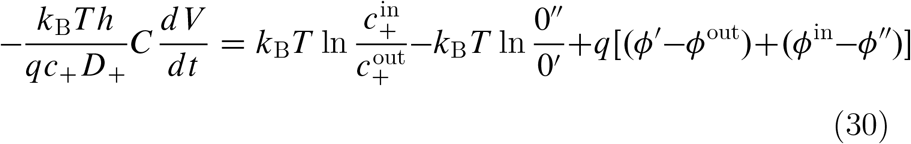

The superscript double prime on the zero in the second logarithmic term indicates the origin of this divergent ln 0 term in the zero concentration of the + ion just inside the membrane at *x* = *h* (to match by electroneutrality the zero concentration of the impermeant anion there). The divergent ln 0 term marked with a single prime has a similar origin at the *x* = 0 surface.

The combined ln(0/0) term is indeterminant at this point in the derivation, but it will turn out to have a sharp physical meaning. The potentials *ϕ* ^″^ and *ϕ* ^′^, respectively just inside the membrane at *x* = *h* and at *x* = 0, are also singular (see Figure 4 in Chapter 3), but the difference *ϕ*^″^ − *ϕ*^′^ will also turn out to have a sharp physical meaning.

To continue, note that the voltage *V* = *ϕ*^in^ − *ϕ*^out^ across the entire energy barrier can be written as the sum of its surface and interior components,

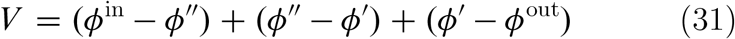

We can then write,

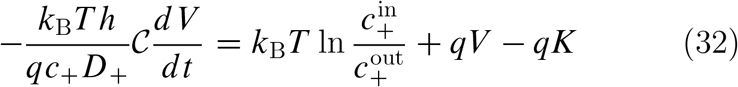

where *qk* stands for the singular terms,

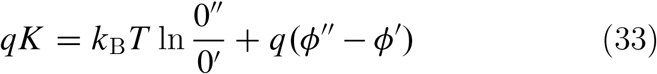

With the definition of the resistance ℛ,

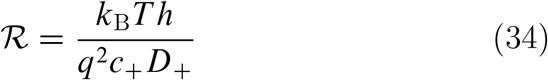

and then on recognizing the Nernst potential for the + ion,

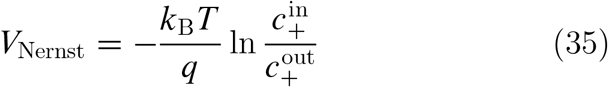

eq ?? becomes,

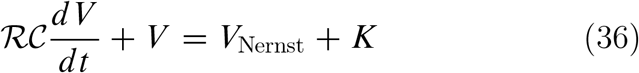

If *k* is assumed constant, the solution to this equation can be written in the form,

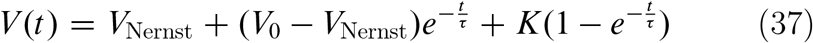

where *V*_0_ is the voltage at *t* = 0, and the time constant *τ* = ℛ 𝒞.

We can look at two special cases. In the first case, the initial voltage *V*_0_ is not equal to *V*_Nernst_, and the system relaxes passively as *t* → ∞ from *V*_0_ to the resting voltage *V*_Nernst_, requiring *k* = 0. It can be checked from eq ?? for *k* and eq 7 in the previous chapter that the equilibrium value of *K* is indeed equal to zero, so the assumption being made in taking *K* as a constant is that the conditions at the boundaries of the energy barrier are equilibrated during the entire time course.

In a second case the system is initially at rest, so that *V*_0_ = *V*_Nernst_, and then at time *t* = 0 a constant current *I* is imposed from an outside source. The voltage then changes from *V*_Nernst_ to *V*_∞_ according to

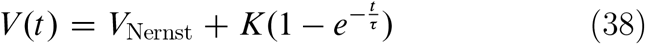

The constant *k* equals *V(*∞) − *V*_Nernst_, which in turn equals *IR*. It is possible to extract a number from these considerations that could be of interest. The resistance ℛ is given by eq ??. It contains the product *D*′*c*′, for which the very small value 1.4×10^−*14*^ mol/(m sec) is obtained from the time constant *τ* = ℛ 𝒞 ≈ 1×10^−*3*^ sec and the capacitance *C* ≈ 0.01 F/m^*2*^ as typically observed of biological membranes. Such a small value is perhaps consistent with an electrical resistance originating at the membrane surfaces, where the diffusion coefficient of the + ion is impeded by a sharp increase *u*′ of entrance energy, and a sharp drop in concentration *c*′ occurs to maintain electroneutrality in the face of the impermeability of the membrane to the anion.

## V. FREE DIFFUSION OF A MIXED SALT

In Chapter II the diffusion constant of a binary electrolyte such as NaCl or KCl was expressed through the Nernst-Hartley relation in terms of the individual diffusion constants of the anion and cation. There is an electric field inside the salt concentration gradient, and the source of the electric field is the polarization charge density due to the different mobilities of the cations and anions.

For biological applications, the minimal electrolyte solution of interest is a mixture of Na^+^, K^+^, and Cl^−^ ions. Discussion of the qualitative difference between the diffusion of binary and mixed salts has a long history, seemingly capped by Robinson and Stokes [13], whose treatise contains theory and data only for the binary salt case, since “where more than two ionic species are present the situation becomes more complex, since there is

an infinite number of ways of satisfying the electroneutrality condition; general equations can be derived, but not necessarily solved for such cases.” Other standard references do not even mention diffusion of mixed salts [14]. Given the centrality of the interplay between sodium and potassium ions in biological electricity [6,7], however, a reassessment of diffusion in mixed salts may be justified. As in Chapter II, there are no membranes or energy barriers in this chapter, just free diffusion of the ions.

The starting point is a set of three electrodiffusion equations, one for the flux *j*_1_ of univalent cations of species 1 (for example, Na^+^), one for univalent cations of species 2 (for example, K^+^), and one for the univalent anion of species designated by a minus sign (for example, Cl^−^),

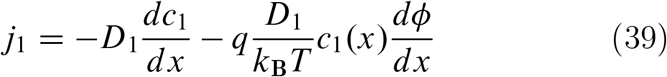

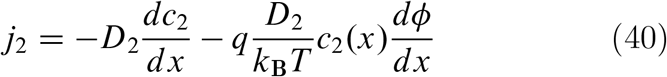

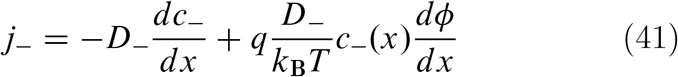

Temporal accumulation of charge is not allowed, since [13] “it is an experimental fact that a macroscopic charge separation does not occur.” Therefore,

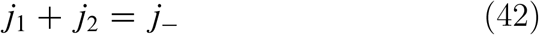

This constraint allows the addition of the equations for *j*_1_ and *j*_*2*_ to result in an independent second equation for *j*_−_,

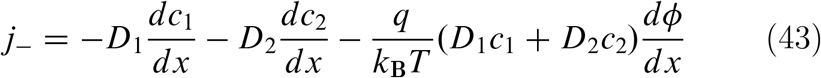

The requirement of local electroneutrality also imposes a constraint on the concentrations,

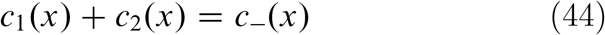

These last two equations suggest a transformation of the variables *c*_1_ and *c*_*2*_ to a pair of new variables *c*_*d*_ and *c*_*s*_,

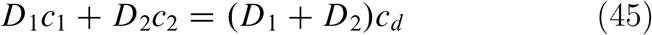

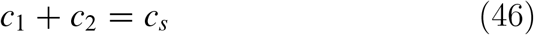

The variable *c*_*d*_ is just an average of the two cation concentrations weighted by their respective diffusion coefficients, while *c*_*s*_ is the total cation concentration in a volume element at *x*. From electroneutrality, *c*_*s*_(*x*) is equal to *c*_−_(*x*). The anion concentration therefore is a measure of the concentration of mixed salt. Analogously, the anion flux *j*_−_ may be identified as the flux *j*_*s*_ of the mixed salt. The cation composition of mixed salt varies throughout its concentration gradient.

The variables *c*_*d*_ and *c*_*s*_ are defined as linear combinations of *c*_1_ and *c*_*2*_. The determinant of coefficients equals *D*_1_ − *D*_*2*_, and therefore the linear transformation is well-defined if the cation diffusion coefficients are not equal. The inverse transformation is obtained by solving for *c*_1_ and *c*_*2*_,

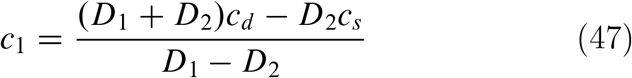

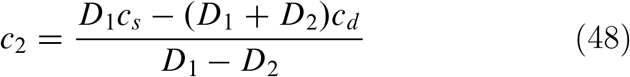

from which the derivatives *dc*_1_/*dx* and *dc*_*2*_/*dx* can be written as needed as linear combinations of *dc*_*s*_/*dx* and *dc*_*d*_ /*dx*.

Return now to the two equations for *j*_−_, writing *j*_*s*_ for *j*_−_ and *c*_*s*_ for *c*_−_, and substituting the new variables and their derivatives,

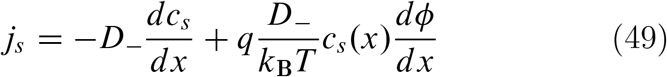

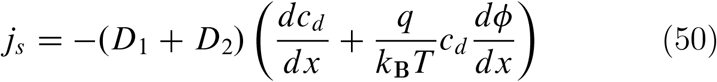

Eliminate *dϕ/dx* by solving for it from the first of these equations,

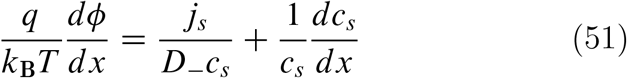

Substitute this expression into the second of the equations for *j*_*s*_, and then solve for *j*_*s*_. The result is,

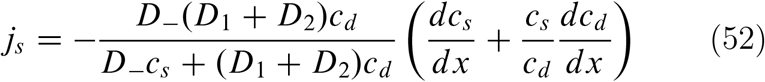

This equation gives the diffusion flux of a mixed salt with two univalent cations and a common univalent anion. The flux of the mixed salt unavoidably depends not only on the salt concentration gradient as in Fick’s law but on the mobility averaged gradient of the two cation concentrations as well.

To get the electric field, or diffusion potential, inside the concentration gradients, substitute this formula for *j*_*s*_ into the previous equation for the potential gradient and simplify,

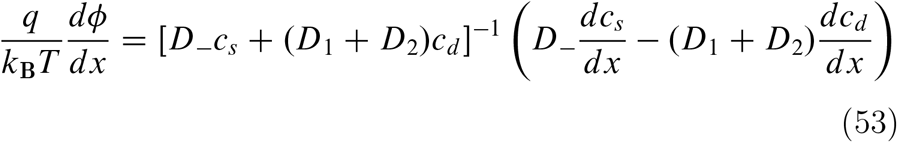

Here is where the difficulty enters. Multiply the preceding equation through by *dx* to obtain a differential form for the potential,

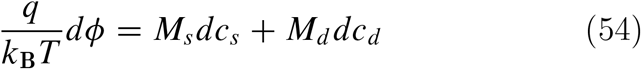

where the coefficients are,

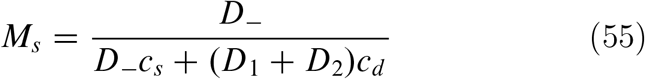

and,

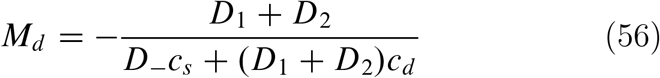

are functions of the variables *c*_*s*_ and *c*_*d*_. It can be checked that the partial derivatives *∂M*_*s*_/*∂c*_*d*_ and *∂M*_*d*_ /*∂c*_*s*_ are not equal (one is the negative of the other), so that *dϕ* is not a perfect differential, and *ϕ* is not a well-defined function. The consequence is stark. The electrodiffusion equations for a mixed salt assume that an electric field operates inside the ion concentration gradients to exert an electric force on the ions. A physically real electric field is the negative gradient of an electric potential *ϕ*.

But the −*dϕ/dx* term in the electrodiffusion equation has no physical meaning because *ϕ* is not a physically meaningful function in the context of these equations, and so has no derivative.

There is a way out of this predicament if one is willing to work with a linearized version of the electrodiffusion equations.

To linearize, it is only necessary to take the ion concentrations as constants, while leaving linearity in the gradients. For example, if we start with a uniform equilibrium solution of NaCl and KCl, and then impose small diffusion gradients, the ion concentrations can be the original uniform ones. The concentrations *c*_*s*_ and *c*_*d*_ appearing above in *M*_*s*_ and *M*_*d*_ are then constants (independent of *x*), and the partial derivatives of *M*_*s*_ and *M*_*d*_ are all equal to zero. The linearized potential *ϕ* is therefore a well-behaved function, and the electric field derived from it gives rise to a physically real force on the ions.

As a final result in this chapter, the equation for the polarization charge density can be given, since the required differentiation of *dϕ/dx* is straightforward once linearization with constant concentrations is done. From the work in Chapter 2 on the diffusion of a binary salt, the free charge density within the concentration gradient is zero by electroneutrality, but the charge density *ρ*_P_ from polarization of the ion distribution that is the source of the electric field is given by − *ϵ*_0_*d* ^*2*^*ϕ/dx*^*2*^ where *ϵ*_0_ is the vacuum permitivity, or, from the present result for *dϕ/dx* with constant coefficients,

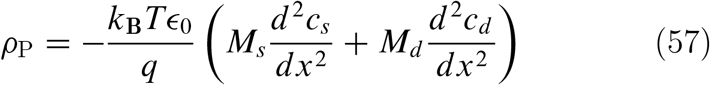

The results in this chapter will be important in the next, but as an interesting example here of how to use these equations, consider the potential arising from placement of a solution of NaCl and another of KCl, of equal concentrations, into direct contact before the two cations have appreciably mixed. In this case *dc*_*s*_, the chloride concentration across the boundary between the two solutions, equals zero, and the expression for *dϕ*, Eq. ??, reduces to *dϕ* = (*RT/* ℱ *)M*_*d*_ *dc*_*d*_. Since only one variable is involved, *ϕ* is a true electrostatic potential, and linearization is not required. In the expressions for *c*_*d*_ and *M*_*d*_, Eqs. ?? and ??, *c*_*s*_ is the constant chloride concentration, and *c*_*2*_ = *c*_*s*_ − *c*_1_ from eq ??. Everything in *dϕ* can be expressed in terms of the single variable *c*_1_, and then *dϕ* can be directly integrated with respect to *c*_1_ across the boundary between the two solutions. If cation 1 is the sodium ion, *c*_1_ = 0 in the potassium solution, and it equals *c*_*s*_ in the sodium solution. Interestingly, *c*_*s*_ winds up having canceled and is not present in the final result,

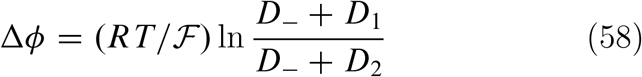

From the known values of the mobilities of the chloride, sodium, and potassium ions *Δϕ* = −*4*.*6mV*, with the sodium solution negative with respect to the potassium solution (since there is greater polarization of the charge distribution on the NaCl side, given the smaller mobility of the sodium ion compared to potassium). Like the equation for the diffusion potential of a binary salt, this formula for *Δϕ* has a long history [1]. In subsequent sections we give numerical examples for more complicated situations involving membrane energy barriers where the problem cannot be reduced to one variable and linearization cannot be avoided.

## VI. THE RESTING POTENTIAL

The preceding chapter was concerned with free diffusion of a mixed salt. For multi-ionic diffusion across a membrane, the interfaces of the membrane with the abutting electrolyte solutions becomes an additional feature that often has not been fully appreciated. Boundary conditions can be trivialized by setting a solute concentration just inside the membrane surface equal to the concentration just outside. A situation for which this boundary condition is obviously inapplicable is ordinary osmosis (solvent flow) induced by an impermeant electrically neutral solute. In this case the solute concentration just outside the membrane drops to zero just inside the membrane. This discontinuity is associated with a discontinuous pressure drop at the membrane surface relative to the value of the pressure in the bulk interior of the solution [10, 11]. A discontinuous pressure drop is the driver of Donnan osmosis also in a system with an impermeant ion together with permeant counterions and added salt composed of permeant ions (see subsequent chapter on Donnan osmosis). In the latter system, as in the former, the concentration of impermeant ion is discontinuous at the membrane, dropping to zero just inside, but the concentrations of the permeant ions are also discontinuous if to enter the membrane requires them to surmount a finite energy barrier (infinite for the impermeant ion). But if the concentrations of these electrically charged particles are discontinuous at the membrane surfaces, one expects the electrostatic potential to be discontinuous at the surfaces also. Introduction of a partition coefficient to represent the ratio of unequal ion concentrations at the membrane surface does nothing toward recognizing the effect on the electrostatic potential. The calculations in this chapter will demonstrate that discontinuous jumps of electrostatic potential are the primary contribution to the resting potential of an energy barrier model for a biological membrane, although the diffusion potential in the interior of the membrane becomes important as well when the energy barriers for the various permeant ions change.

Figure 5 illustrates the discontinuities at the membrane surfaces of all quantities in a Donnan ionic system, pressure, potential, and ionic concentrations. The impermeant anion plays no role in the calculations of this chapter but is important in the next one. Figure 6 shows the energy barrier model with different levels of mechanical energy for different ions.

**Figure 5.**
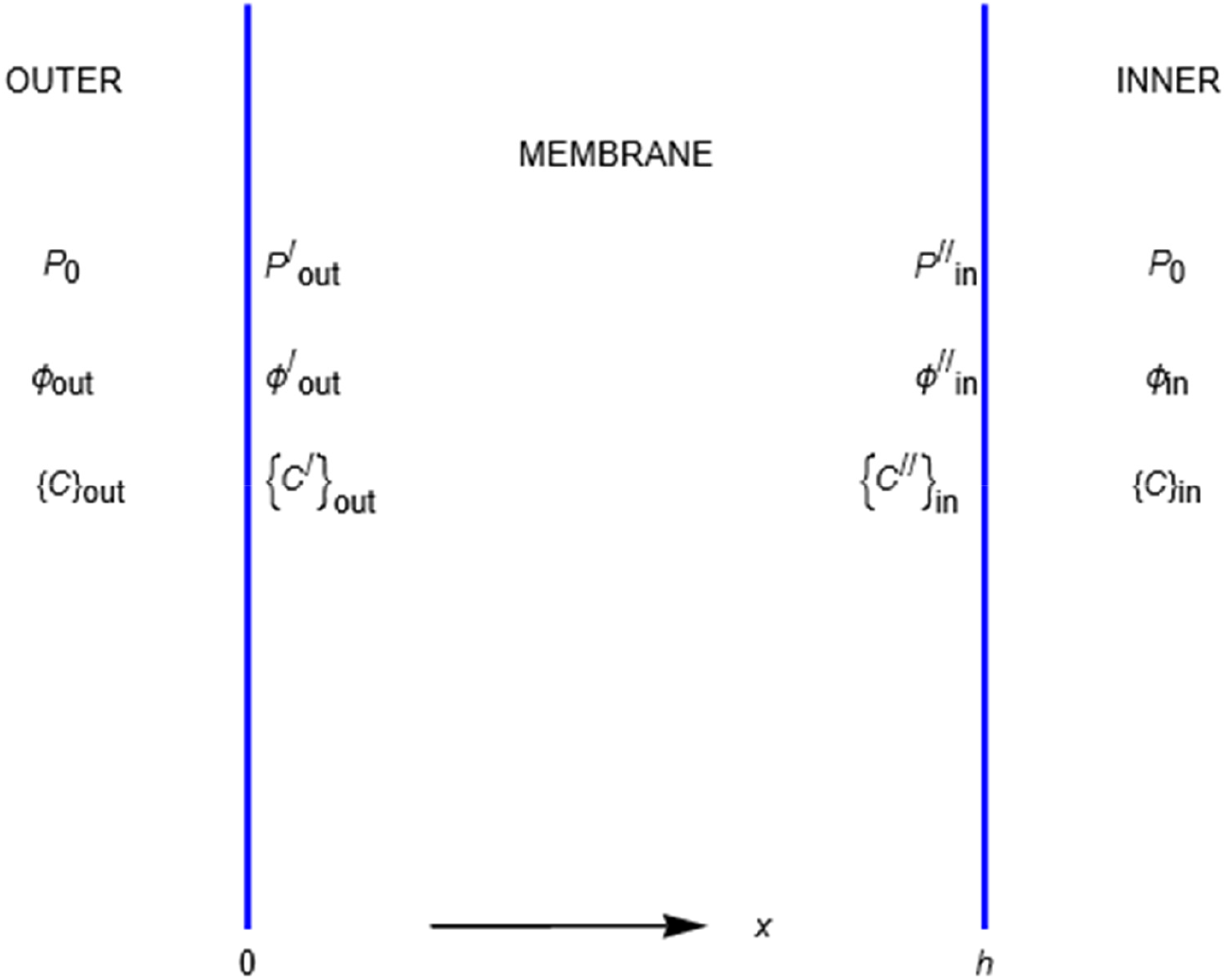
The complete Donnan ionic system, showing the relevant quantities pressure, electric potential, a set of small ion concentrations {c} in the outer compartment x<0, and a set of ion concentrations in the inner compartment, x>h, which includes the concentration of an impermeant anion. These quantities are discontinuous at the membrane boundaries x=0 and x=h, and their values just inside the membrane are designated by primes. The value of an impermeant anion concentration just inside the membrane is zero, but the concentrations of permeant ions inside the membrane are not zero.

**Figure 6.**
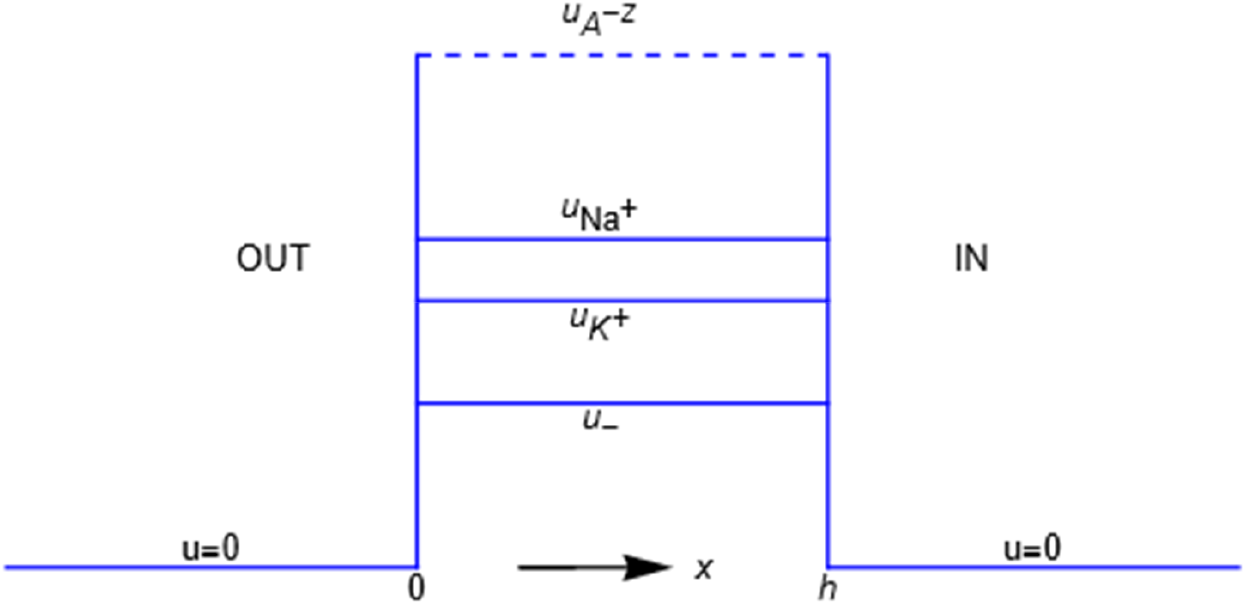
Profiles of the Debye potential energies u_i_ representing the mechanical interactions of the membrane with the four types of ions considered in this paper. The ordering of the potential energies of the small ions is arbitrary. The impermeant anion A^-z^ is at *u*=∞.

The electrostatic potential difference between the inner and outer electrolyte solutions bathing the membrane (energy barrier) that separates them can be analyzed as three distinct parts,

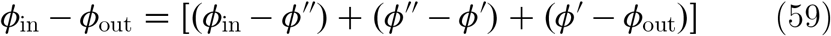

where the first term on the right-hand side is the jump at the membrane surface between the membrane interior and the inner compartment, the last term is the jump at the membrane surface between the membrane interior and the outer compartment, and the central term is the internal potential difference between the two membrane sides. Note how two obvious cancellations on the right-hand side show that the right-hand side is equal to the overall potential difference *ϕ*_in_ − *ϕ*_out_.

To emphasize the importance of the jumps of electrostatic potential at the membrane surfaces, the study begins with them and only subsequently considers the interior of the membrane. As in the previous chapter on free diffusion of mixed salts, univalent cation 1 here could be Na^+^, univalent cation 2 could be K^+^, and the univalent anion designated “−”, could be the Cl^−^ ion. The Donnan impermeant anion is not involved in the calculation of the resting potential. The flux equations contain the interface ion-membrane mechanical forces, but we omit solvent drag (bulk flow) terms, which can be included (and are in the next chapter on Donnan osmosis) but would disappear from the calculation here and have no effect on the results.

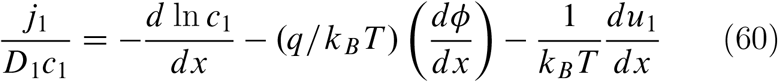

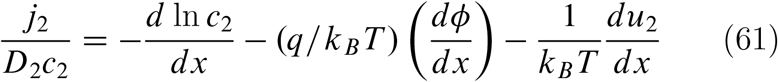

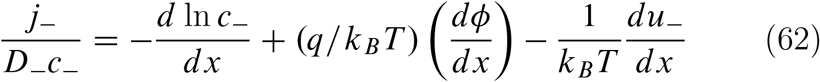

Add eqs ?? and ??,

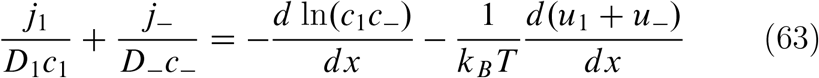

Add eqs ?? and ??

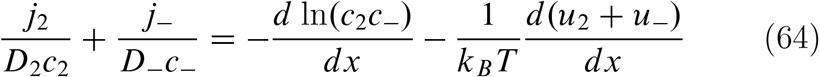

Integrate eq ?? across the left interface in Figure 5, left to right (increasing *x*), noting that integration of the left-hand side gives zero, since the width of the interface is zero,

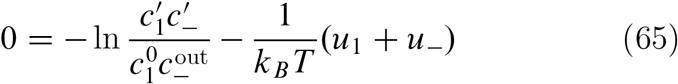

The same integration of eq ?? gives,

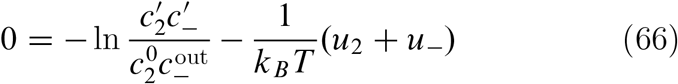

The next pair of equations is simply a respective rearrangement of the above two, with the definitions *α*_*i*_ = exp(−*u*_*i*_*/k*_*B*_*T), i* = *1, 2*, −,

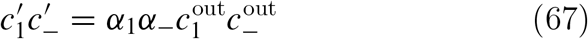

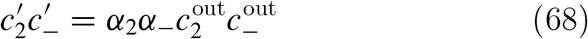

Electroneutrality provides a third equation for the three unknowns 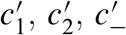,

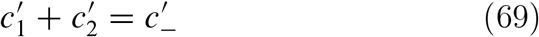

Solving,

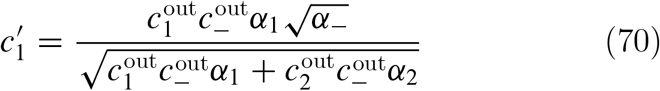

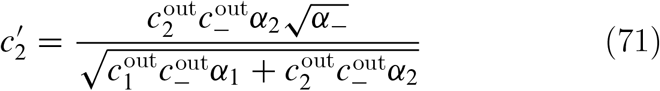

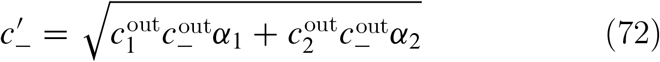

It is important to notice that these ion concentrations just inside the membrane have a complicated dependence both on the concentrations outside the membrane and on the energy barrier parameters *α*_*i*_. For example, 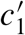 is *not* equal to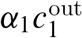. In fact, the concentration of cation 1 just inside the membrane depends not only on its concentration outside, but it also depends on the outside concentrations both of cation 2 and the anion, as well as on the barrier energy for cation 2.

To get the potential jump at the left interface, integrate eq ?? across this interface, left to right, from outside to just inside the membrane,

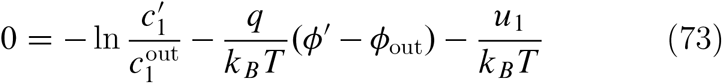

Substitute the expression for 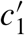 from eq ??, and the potential jump then follows from a bit of algebra and cancellation,

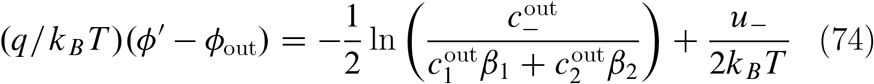

This expression gives the value of the potential just inside the membrane minus its value outside at the membrane surface between the outer electrolyte solution and the membrane.

The calculation for the potential jump at the right interface follows exactly the same procedure as at the left, but two of the equations can be recorded, because the integrations from left to right (increasing *x*) are from just inside the membrane to outside (see Figure 5), and careful attention must be paid to the signs.

Integration of eqs ?? and ?? across the right interface gives,

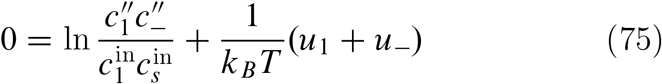

and,

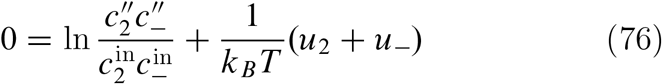

The equations for the concentrations 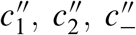 just inside the membrane at the right interface are the same as eqs ??, ??, and ?? with 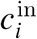 replacing 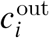. Finally, the potential jump at the right interface is obtained by integrating eq ?? across the right interface from just inside the membrane to outside,

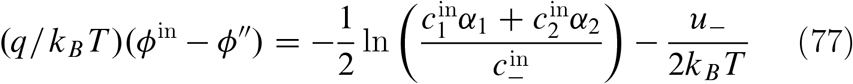

an expression for the value of the potential outside the membrane minus its value just inside at the membrane surface between the inner electrolyte solution and the membrane.

Let *Δϕ*_boundary_ be the contribution of the two surface jumps to the total potential across the membrane. It is the sum of eq ?? and eq ??,

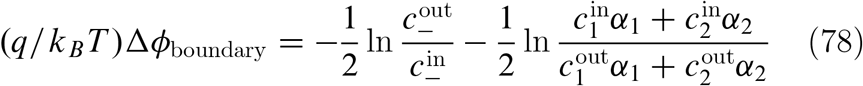

The left-hand side is the boundary contribution to the voltage *V*_m_ across the membrane, rendered dimensionless by multiplying it by *q/k*_*B*_ *T*, which equals ℱ */RT*, where ℱis the Faraday and *R* the gas constant. The first term on the right is just half the Nernst potential 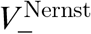 for the anion.

As a numerical example consider a sodium-potassium-chloride system in a physiological context when a cell membrane is at rest [3, 6]]. Typically, the Na^+^ ion (cation 1) is much less permeant than the K^+^ ion (cation 2), so set *α* _1_ = 0, obtaining for the boundary contribution,

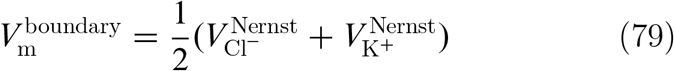

Using typical values for the Nernst potentials of animal cells, 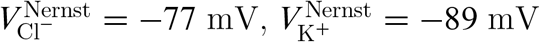, results in the value −*83* mV for the boundary contribution, roughly typical of resting potentials for animal cells. On the other hand, if *α* _*2*_ were to be set to zero, mimicking a shutdown of the ability of the K^+^ ion to enter the membrane, then

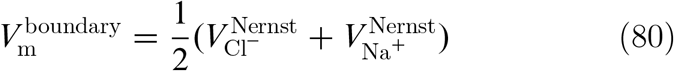

and with 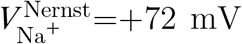, the boundary contribution to the membrane potential becomes depolarized to the value −3 mV.

Turn now to the potential change across the interior of the membrane, that is, the second term in eq ??, *ϕ* ^″^ − *ϕ* ^′^. For it, use the results of the previous chapter for the diffusion potential of a mixed salt. Recall that the ion concentrations in the coefficients *M*_*s*_ and *M*_*d*_ are constant, and take them here for numerical purposes as averages; for example, in the interior of the membrane 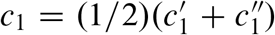. The primed concentrations are on the membrane side of the discontinuities and are calculated from the external concentrations. For them, typical values in the cytoplasm and extracellular fluid of an animal cell can be found in [3].

For the case of impermeant Na^+^, *α* _1_ = 0, take the potassium and chloride ions as freely permeant, *α* _*2*_ = *α* _−_ = 1. Assume as well that the diffusion coefficient for the sodium ion inside the membrane is zero, to be consistent with the assumption that it is impermeant. The diffusion coefficients for the K^+^ and Cl^−^ ions are set equal to their values in water. Then the diffusion potential, that is, the voltage change across the interior of the membrane, numerically computes as equal to 0.11 mV, insignificant compared to the boundary voltage −83 mV found above. On the other hand, for the case of impermeant potassium and chloride ions but freely permeant sodium, the contribution from the diffusion voltage is very large, 53 mV, so the overall voltage across the membrane is +50 mV. This behavior is reminiscent of the changes occurring during the action potential.

## VII. OSMOSIS

In the two compartments separated by a semipermeable membrane that illustrate osmotic pressure, one compartment contains pure water, the other an aqueous solution of a single neutral solute (like sugar), and the membrane is permeable to water but impermeant to the solute. Then at equilibrium the pressure in the solution is higher than in the pure solvent, and the pressure difference is called the osmotic pressure. If the pressure is constrained to be equal in both compartments, then the steady flow of water from the compartment with pure water into the compartment containing the solution is called osmosis. Both osmosis and osmotic pressure are now understood mechanistically to arise from an abrupt pressure drop at the membrane-solution interface [8–11]. When the solute is an overall electroneutral electrolyte like NaCl, and one but not both of the dissociated ionic components of the electrolyte are impermeant, the permeant ion nevertheless cannot escape the solution because an infinite electrostatic potential develops that enforces electroneutrality of the ionic solution (see the introductary chapter).

The situation in a Donnan system is different. In a prototype Donnan system, a semipermeable membrane again separates two compartments. In one compartment there is a salt solution like aqueous NaCl, and both the sodium and chloride ions are permeant to the membrane. In the other compartment resides an impermeant anion (usually a large one, such as a protein) along with the sodium and chloride ions. At equilibrium a pressure difference called Donnan osmotic pressure has developed. But also there is an equilibrium Donnan electrostatic potential across the membrane. Moreover, there is an equilibrated redistribution of the small ions such that the chloride ion is partially excluded from the compartment containing the impermeant anion, sometimes called Donnan salt exclusion. The thermodynamic analysis of all three interconnected Donnan equilibrium effects is not as easy to find as it should be, but the equations for all three effects in ideal dilute solutions are available here in the Appendix.

The Donnan counterpart to osmosis is a steady bulk volume flow into the solution containing the impermeant ion that develops when the pressures in both compartments are constrained to be equal. The volume flow may aptly be called Donnan osmosis. Donnan osmosis is caused on a mechanistic level by abrupt and unequal pressure drops at the two surfaces of the membrane. It follows, as in ordinary osmosis, that there exists a pressure gradient inside the membrane from one side of the membrane to the other. It is this interior pressure gradient that drives water flow across the membrane. The result for the equation governing Donnan osmotic volume flow across an energy barrier is presented and discussed first. The step-by-step derivation follows.

The calculation is for the mimimal system of interest in biology, an impermeant anion A^−*z*^ with charge −*zq* in a mixture of permeant Na^+^, K^+^, and Cl^−^ ions (see Figure 5 in the previous chapter). The result for the Donnan volume flow *J*_*v*_ per unit area is,

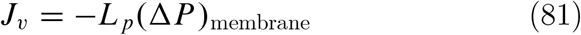

where (Δ*P*)_membrane_ is the internal pressure difference between the two sides of the membrane, and the coefficient *L*_*p*_ is the hydraulic permeability. The expression for (Δ*P*)_membrane_ is,

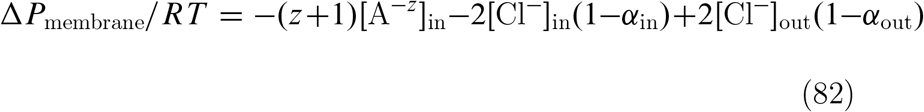

where *R* is the gas constant, *T* the Kelvin temperature, and the bracketed quantities are concentrations in molarities.

The first term on the right side of the equation for Δ*P*_membrane_ should be familiar. It is the sum of a van’t Hoff term [A^−z^]_in_ and another van’t Hoff term *z[*A^−z^]_in_ accounting for the *z* univalent sodium and potassium counterions required to remain in the inner compartment to neutralize (electrostatically) the charge −*zq* on the impermeant anion A^−z^. Notice that it makes no sense to try to specify what proportion of the neutralizing counterions are sodium and what potassium. The neutralization is longrange electrostatic in nature, and all that can be said is that *z* among the sodium and potassium ions are rendered effectively impermeant by each mechanically impermeant A^−z^ anion.

The two further terms on the right side of the equation contain the effect of the permeant Na^+^, K^+^, and Cl^−^ ions other than those required to neutralize the charge on the A^−*z*^ anion.

Only the concentrations of the chloride ions appear explicitly in these electrostatic terms because the combined charge of these “other” sodium and potassium ions is constrained to neutralize the charge on the chloride ions, accounting for the factor 2. The concentrations of sodium and potassium ions do appear in the definitions of the quantities *α*_in_ and *α*_out_,

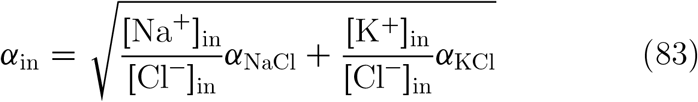

and,

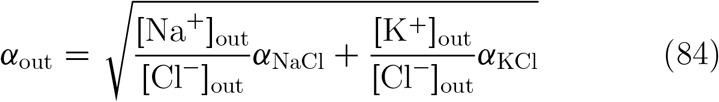

In these expressions, *a*_NaCl_ and *a*_KCl_ are defined as,

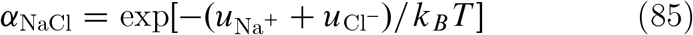

and,

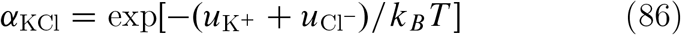

In these last two equations, 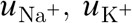, and 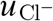 are the energy barrier heights for the indicated ions, and *k*_*B*_ is the Boltzmann constant (gas constant *R* divided by Avogadro’s number).

A useful observation follows if the membrane energy barriers are actually barriers, in other words, if all of the energies *u*_*i*_ are positive. Then the quantities *α*_NaCl_ and *α*_KCl_ are less than unity, and it is easy to show from the electroneutrality conditions that the concentration-averaged quantities *α*_in_ and *α*_out_ defined in Eq. ?? and Eq. ?? obey the inequalities 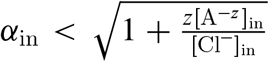 and *a*_out_ *< 1*. If all of the barrier heights *u*_*i*_ are zero, so that there are no membrane impediments to transit of the sodium, potassium, or chloride ions, the inequalities become equalities,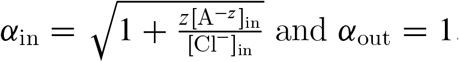 and *α*_out_ = 1.

Two special cases are worth making explicit, Donnan osmosis if the small ions are completely impermeant along with the impermeant anion A^−*z*^, and Donnan osmosis if the small ions are freely permeant. For impermeant sodium, potassium, and chloride ions, the corresponding Debye energy barrier levels *u*_*i*_ are infinite, so that *α*_in_ = *α*_out_ = 0. The expression for volume flow reduces to,

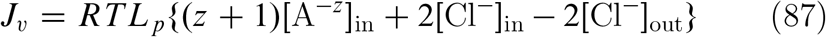

which would be the same as ordinary osmosis when all solutes are impermeant. If at the opposite extreme the small ions are freely permeant, the barrier energies for them are all zero, and Donnan volume flow becomes,

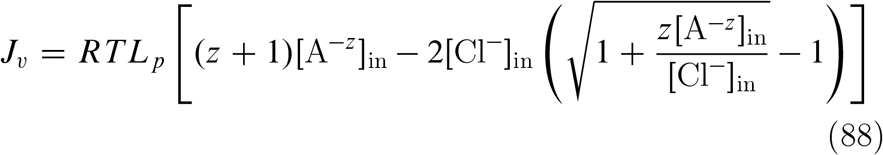

In this case the small ions exert an effect on osmosis even though they are mechanically freely permeant, indicating the important influence of the Donnan electrostatic potential.

Figure 7 shows a 3D plot of Δ*P*_membrane_*/RT*[A^−*z*^]_in_. The impermeant anion A^−*z*^ is present at concentration 0.144 M. Ion concentrations for a typical animal cell are [Cl^−^]_in_=0.006 M, [Cl^−^]_out_=0.106 M, [Na^+^]_in_=0.010 M, [K^+^]_in_=0.140 M, [Na^+^]_out_=0.145 M, [K^+^]_out_ = 0.*005* M. The combined sodium and potassium concentrations are equal in the extracellular fluid (“out”) and the intracellular cytoplasm (“in”), but Na^+^ is enriched outside the cell, and K^+^ inside. The concentration of impermeant anion has been chosen to meet the requirement of electroneutrality for *z*=1. Figure 7 makes plain that the values of Δ*P*_membrane_ are negative nearly everywhere, driving osmotic flow *J*_*v*_ into the inner compartment containing the impermeant anion. Only along the axis *α*_NaCl_ = 0 does *J*_*v*_ vanish, whereas along this axis the value of *a*_KCl_ ranges from zero (impermeant) to unity (freely permeant).

**Figure 7.**
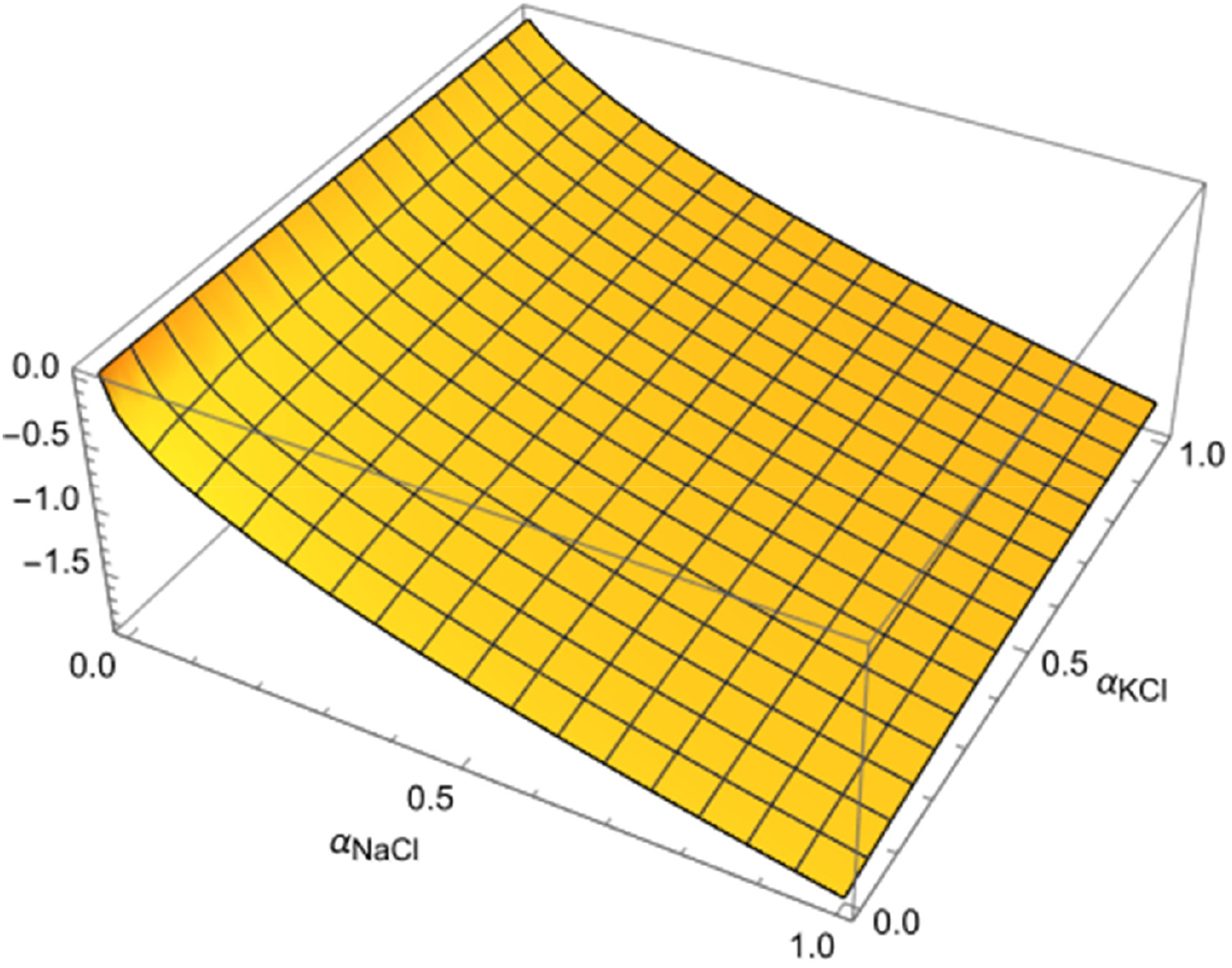
The surface is the negative of the normalized osmotic volume flow J_v_ for ionic concentrations typical of animal cells. The negative values imply positive osmotic water flow into the inner compartment. The values are near zero where α_NaCl_=0, i.e., impermeant Na+ ions. These values are near zero for any value of α_KCl_. This behavior is consistent with the action of the sodium ion pump.

The results in Figure 7 are consistent with an informal interpretation of the action of the sodium pump [3]: “… Na^+^ ions, the most abundant permeant cations outside the cell, are extruded from the cell by active transport as rapidly as they leak in … functionally equivalent to making the cell membrane impermeable to Na^+^ ions”, and thus “forestalling osmotic catastrophe” (swelling and ultimate bursting of the cell). Indeed, the graph in Figure 7 shows that osmotic water flow into the cell is reduced to zero or near-zero values when *α*_NaCl_ = 0, that is, when the Na^+^ ion is impermeant.

As another example, the implication of Donnan osmosis for transport across capillary walls may be of interest. Capillary walls are highly porous, and small solutes, including ions, pass through easily [3]. The special case Eq. ?? for freely permeant ions is therefore applicable. With the relevant concentrations specified by Blaustein et al. [3], and the value *z* = 1 [19], the value of *J*_*v*_*/RTL*_*p*_ equals 0.82 (this flow from osmosis does not include the flow caused by the hydrostatic blood pressure). Of possibly greater interest than the number itself is that the volume flow would have been much greater if the negative contribution of the second term in Eq. ?? had been absent. This term is precisely what distinguishes Donnan osmosis in this case from the more familiar van’t Hoff osmosis. It accounts for the electrical effect of the permeant small ions, which is not taken into account in the usual listings of “Starling forces” responsible for capillary transport. Similar considerations may be applicable to other porous physiological structures.

### Derivation

In the derivation the notation for concentrations has been streamlined. The concentrations of the sodium and potassium ions are respectively *c*_1_ and *c*_*2*_, the concentration of the chloride ion is *c*_−_, and the concentration of the impermeable anion is *c*_*a*_. The units of concentration are number of molecules per unit volume, so that Boltzmann’s constant *k*_*B*_ replaces the gas constant *R*.

In the theory of the hydrodynamics of fluids, the fundamental equation for uniform volume flow in a direction *x* expresses the imbalance of forces on a volume element of fluid,

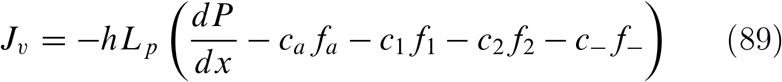

For flow across a membrane of thickness *h*, the coefficient *L*_*p*_ is the hydraulic permeability. The collection of terms in parentheses is the total force acting on a cross-sectional slab of width *dx* located at *x*. The pressure derivative is the net force exerted per unit area on the two planar surfaces of the slab. The second term is the mechanical force *f*_*a*_ from the membrane material exerted on molecules of the impermeant anion located within the slab.

Of course the concentration *c*_*a*_ of the impermeant anion is zero inside the membrane, and the repelling force *f*_*a*_ acts only at the membrane interface with the inner compartment containing it (*x* = *h* in Fig. 5). The following three terms are the mechanical forces on the sodium, potassium, and chloride ions, and these forces also are nonzero only at the interfaces, because the environment of an ion inside the membrane (the energy barrier as modeled here) is symmetric on both sides of it. The forces *f*_*i*_ made explicit in this equation are the mechanical ion-membrane forces. There are also electric forces on all of the ionic charges due to an electric field −*dϕ/dx*. But the slab is net electrically neutral, as the total charge of all the ions inside it sum to zero, so the total electric force on the slab is zero.

In this equation the forces *f*_*i*_ could have been expressed as in previous chapters as negative gradients of potential energies, but the direct force notation emphasizes that the pressure *P* is also a mechanical force per unit cross-sectional area, and that osmosis is driven by an imbalance of mechanical forces. The derivation then proceeds through the equations for the fluxes *j*_*i*_ of the various species including the effect of the electrostatic potential. The flux *j*_*a*_ of the impermeant anion is set to zero, and use of electroneutrality is a central condition both outside and inside the energy barrier.

The beginning step is the elimination of the mechanical force *f*_*a*_ on the impermeant anion. The equation for the flux *j*_*a*_ of this ion is,

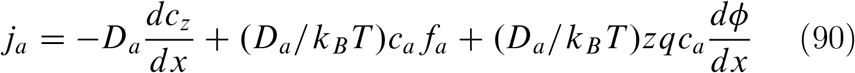

where *D*_*a*_ is a diffusion constant. But *j*_*a*_ = 0 since the A^−z^ ion is impermeant, and the resulting equation can be solved for *c*_*a*_*f*_*a*_, which can then be substituted back into Eq. ?? to give,

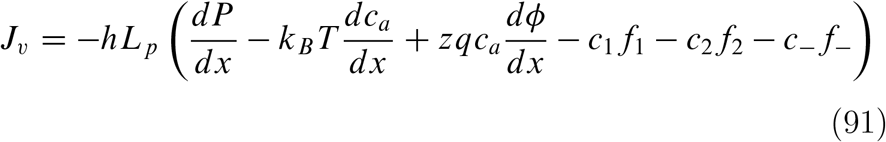

Next, electroneutrality is invoked, *zc*_*a*_ = *c*_1_ + *c*_*2*_ − *c*_−_, in the electric field term,

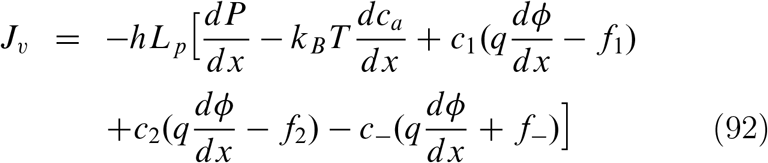

The electrical and mechanical forces on the sodium, potassium, and chloride ions can be eliminated by looking at the equations for their fluxes, which do not vanish since these ions are permeant. For example,

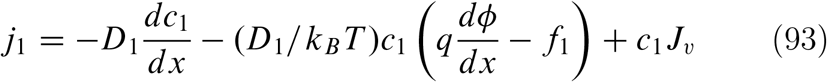

where a solvent drag term has been added (it does not affect the final result). This equation can be solved for the combined electrical and mechanical force term for the sodium ion, which is then substituted into Eq. ??. After analogous substitutions for the potassium and chloride ions, Eq. ?? becomes,

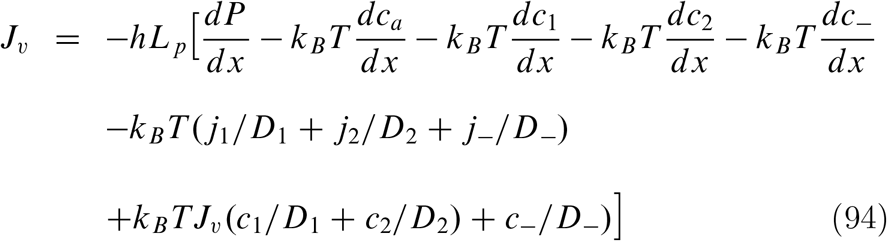

Both sides of this equation can be integrated across the surface separating the outer compartment from the interior of the membrane. The terms involving *J*_*v*_ and the *j*_*i*_ integrate to zero, because the integration is over an infinitesimal interval (just outside the membrane to just inside). The integral of *dP*/*dx* equals the difference 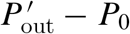, that is, the pressure drop at the outer surface. The integral of the *dc*_*a*_/*dx* term vanishes, because *c*_*a*_ is zero both in the outer compartment and inside the membrane.

As for the permeant ion terms, as an example, the integral of the derivative *dc*_1_/*dx* across the outer interface equals the difference 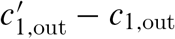, where the first term is the concentration just inside the interface, and the second is the concentration just outside. The result of the integration is,

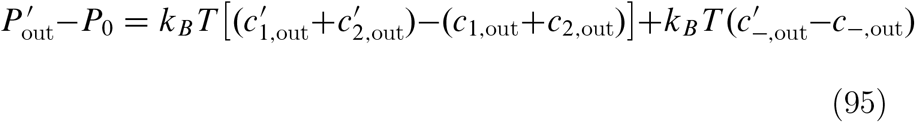

or, from electroneutrality *c*_1_ + *c*_*2*_ = *c*_−_ for both the primed and unprimed concentrations,

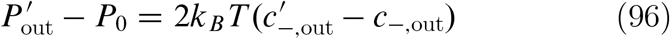

Performing an analogous integration of Eq. ?? from left to right across the inner membrane surface, with electroneutrality here reading *c*_1_ + *c*_*2*_ = *zc*_*a*_ + *c*_−_ outside the membrane in the inner compartment, and 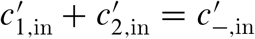 just inside the membrane at the inner surface, results in,

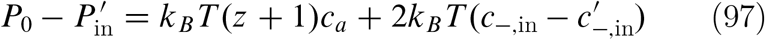

The pressure gradient inside the membrane, from one side to the other, is (Δ*P*)_membrane_, equal to 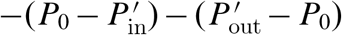, so adding the negatives of the previous two equations,

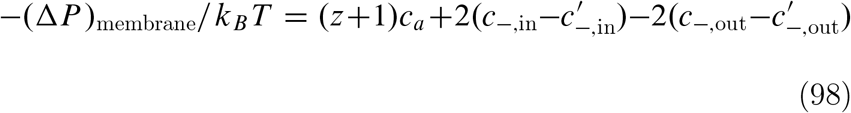

The primed chloride concentrations in this equation for (Δ*P*)_membrane_ are the concentrations just inside the membrane at the interfaces. They are not known at this point in the derivation, and the next task is to find them. Actually, they can be found in the previous chapter, but the derivation is repeated here for completeness. Write the flux equations for the sodium and chloride ions, and add them to eliminate the electric field terms,

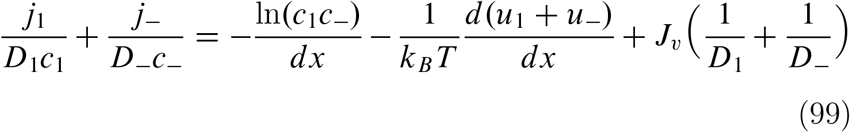

In this equation we have used *f*_1_ = −*du*_1_/*dx* and *f*_−_ = −*du*_−_/*dx*.

Similarly,

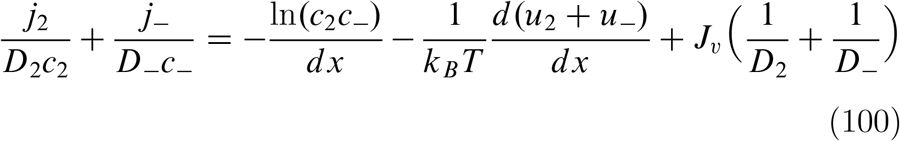

Now integrate both sides of Eq. ?? across the outer membrane surface, noting that *u*_1_ = *u*_−_ = 0 in the outer compartment, and, as above, that integration of the flow terms vanish. After exponentiating,

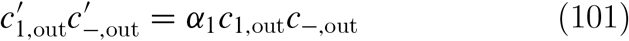

where *α*_1_ is the same quantity as *α*_NaCl_ in Eq. ??. Analogous handling of Eq. ?? yields,

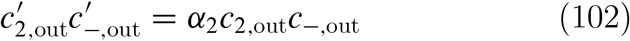

where *α*_*2*_ is the same quantity as *α*_KCl_ in Eq. ??. Together with electroneutrality, 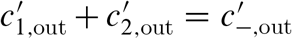, the preceding two equations give three equations for the three unknowns 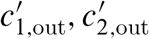 and 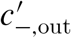, namely the sodium, potassium, and chloride ion concentrations just inside the membrane at its outer surface. Solving,

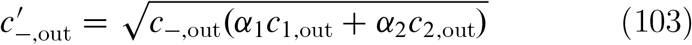

There are expressions for the sodium and potassium concentrations also, but they are not needed here. The inner surface can be handled analogously on noticing that the electroneutrality condition inside the membrane at the inner surface has the same form as for the outer since *c*_*a*_ = 0 inside the membrane.

For the inner surface then,

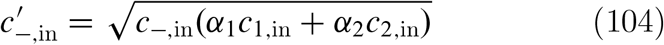

Substitution of these two equations into Eq. ?? results in Eq. ?? and completes the derivation.

## APPENDIX

It is remarkably difficult to find a comprehensive treatment of Donnan equilibrium. Overbeek [20] is a good source, marred only by mystification of the Donnan potential. Expressions for the three equilibrium Donnan effects in ideal dilute conditions are listed here. All of them may be derived either mechanistically along the lines of the text, or by standard thermodynamic methods. The Donnan system consists of a semipermeable membrane separating an aqueous salt solution like NaCl in an “outer” compartment from a corresponding solution in an “inner” compartment that also contains an impermeant anion Z with charge −*zq, z* a positive number, *q* the charge on a proton, and concentration *c*_*z*_. The sodium and chloride ions are permeant. Equilibrium is established between inner and outer compartments. Concentrations in the outer compartment are marked with a “prime”. Concentrations in the inner compartment are unmarked. The salt concentration in the outer compartment is 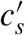 (equal to the sodium and chloride ion concentrations there), assumed known. The salt concentration in

the inner compartment is *c*_*s*_ (equal to the chloride concentration there, the sodium concentration being equal to *zc*_*z*_ + *c*_*s*_).

Define 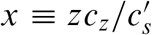 (note the different meaning of *x* here than in the text). Define 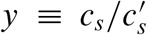 as the Donnan salt exclusion ratio (Donnan ion distribution). Then at equilibrium, *y(x*) is the function of *x* given by the solution of the quadratic equation (*x* + *y)y* = 1.

The Donnan electrostatic potential difference *ϕ* − *ϕ*′ between inner and outer compartments is given by,

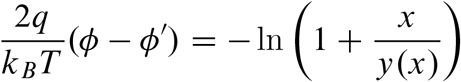

The Donnan osmotic pressure is given by,

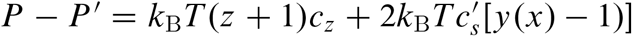

